# Ultrasound Neuromodulation and Correlation Change in the Rat Somatosensory Cortex

**DOI:** 10.1101/2022.03.18.484914

**Authors:** Sandhya Ramachandran, Xiaodan Niu, Kai Yu, Bin He

**Affiliations:** Department of Biomedical Engineering, Carnegie Mellon University; Neuroscience Institute, Carnegie Mellon University

**Keywords:** Neuromodulation, Transcranial focused ultrasound, tFUS, Frequency dependent plasticity

## Abstract

Transcranial focused ultrasound (tFUS) is a neuromodulation technique which has been the focus of increasing interest for noninvasive brain stimulation with high spatial specificity. Its ability to excite and inhibit neural circuits as well as to modulate perception and behavior has been demonstrated, however, we currently lack understanding of how tFUS modulates the ways neurons interact with each other. This understanding would help explain tFUS’s mechanism of high-level neuromodulation and allow future development of therapies for neurological disorders. In this study we investigate how tFUS modulates neural interaction and response to peripheral electrical limb stimulation through intracranial multi-electrode recordings in the rat somatosensory cortex. We deliver ultrasound in a pulsed pattern to attempt to induce frequency dependent plasticity in a manner similar to that found following electrical stimulation. We show that neural firing in response to peripheral electrical stimulation is increased after ultrasound stimulation at all frequencies, showing tFUS induced excitation in individual neurons *in vivo*. We demonstrate tFUS frequency dependent pairwise correlation changes between neurons, with both potentiation and depression observed at different frequencies. These results extend previous research showing tFUS to be capable of inducing synaptic depression and demonstrate its ability to modulate network dynamics as a whole.

## Introduction

Neuromodulation, the use of physical stimulus to modulate information processing and transmission in the nervous system, has gained widespread attention to offer non-pharmacological options to treat neurological diseases such as depression, chronic pain, Alzheimer’s disease, Parkinson’s disease, and epilepsy. One technique which has gained increasing interest is Transcranial Focused Ultrasound (tFUS), the use of a low-intensity acoustic wave to stimulate the brain with acoustic energy with high precision targeting capabilities (Blackmore et al., 2019; Kamimura et al., 2020; Naor et al., 2016; Tufail et al., 2010, 2011; Tyler et al., 2008). Although many other neuromodulatory techniques exist, including deep brain stimulation (DBS) (Ashkan et al., 2017; Gardner, 2013; Lozano et al., 2019), transcranial magnetic stimulation (TMS) (Chail et al., 2018; Koponen & Peterchev, 2020), transcranial direct current stimulation (tDCS) (Bikson & Rahman, 2013; Giordano et al., 2017; Nitsche & Paulus, 2000), and optogenetics (Boyden et al., 2005; Mahmoudi et al., 2017), tFUS has the potential to offer high spatial focality and deep brain penetration as a noninvasive neuromodulation technique applicable to human subjects. Chemical methods and optogenetics can allow for high temporal resolution and/or cell type specificity, but are invasive approaches that require introducing foreign substances to the brain (Chen et al., 2018; Mahmoudi et al., 2017). DBS has shown much promise in treating disorders in humans, especially Parkinson’s Disease, but requires an invasive surgical procedure to insert electrodes deep into the brain for electrical stimulation (Lozano et al., 2019). TMS and tDCS are noninvasive, but come with certain limitations in their spatial resolution as well as deep brain penetration (Fini & Tyler, 2017; Kubanek et al., 2018; Sanguinetti et al., 2020). tFUS stands out as a potential next-generation noninvasive neuromodulation technique with high spatial resolution (1mm resolution) and ability to target deep brain regions (Legon et al., 2014; Mehić et al., 2014), as well as cell type specificity (Legon et al., 2014; Mehić et al., 2014; Yu, Niu, et al., 2021).

tFUS has been shown to elicit action potentials in various *in vivo* and *in vitro* studies in cell culture and brain slices (Muratore et al., 2009; Tyler et al., 2008) as well as across a variety of animals including worms (Kubanek et al., 2018), rodents (Juan et al., 2014; Mehić et al., 2014; Niu et al., 2018; Tufail et al., 2010; Younan et al., 2013; Yu et al., 2016), monkeys (Deffieux et al., 2013; Folloni et al., 2019), as well as humans (Hameroff et al., 2013; Lee et al., 2015; Legon et al., 2014; Liu et al., 2021; Verhagen et al., 2019; Yu, Liu, et al., 2021; Zhang et al., 2021). Effects from stimulation have been found on various many levels recordings including in multi-unit activity (MUA) (Tufail et al., 2010), local field potential (LFP) (Yuan et al., 2015), the blood oxygen level dependent (BOLD) signal (Verhagen et al., 2019), and EEG (Lee et al., 2015; Legon et al., 2014). These effects have been shown to be due to direct activation, not merely indirect activation through an auditory pathway due to some sound from the ultrasound wave (Mohammadjavadi et al., 2019). Ultrasound has also been shown to be capable of causing both excitatory and suppressive neural effects, with the direction of modulation depending on ultrasound frequency and delivery pattern (H. C. Kim et al., 2021; King et al., 2013; Lee et al., 2015). These abilities show that ultrasound is capable of modulating neural function in some way, but the potential effects of this kind of stimulation require further study to determine whether tFUS can function as a therapeutic technique for neurological disorders.

To understand tFUS’s capabilities with regards to treating neurological disorders, it is important to understand its effect on the way that neurons interact with each other in a network. Abnormalities in the ways that neurons fire in relation to each other and overall behave as a circuit are often a marker of pathology. Epilepsy is characterized by unusual synchronized firing, resulting in abnormal high correlation between neurons (Staba et al., 2002; Warren et al., 2010). Parkinson’s disease on the other hand is characterized by synchronized oscillatory behavior in the beta band (15-30Hz) in the subthalamic nucleus (Weinberger et al., 2006). Additionally, use of neuromodulation to alter this type of pathologic behavior has been shown to treat the related disorders. DBS has been observed to attenuate oscillatory activity such as that characteristic in Parkinson’s disease (Weinberger et al., 2006), as well as other abnormal firing patterns and correlations (Daneshzand et al., 2017; Larrivee, 2018; Lozano et al., 2019). It has also been shown to restore connectivity and ability to grow new connections in the hippocampus, resulting in long term restoration of neural learning (Hao et al., 2015; Kai Tan et al., 2020). fMRI (Horn et al., 2019) and DTI (van Hartevelt et al., 2015) studies have investigated the effect of DBS on the functional and structural connectome respectively and shown that DBS can normalize these measurements in patients with Parkinson’s disease while significantly reducing symptoms. One mechanism heavily implicated in this type of connectivity change is synaptic long term potentiation and depression (LTP and LTD).

Electrical stimulation is known to be capable of causing changes in synaptic connectivity through both tetanic stimulation and paired pulse facilitation. Studies have been performed on LTP in the hippocampus, demonstrating that high frequency (50 Hz and higher) tetanic electrical stimulation to Schaffer collaterals causes an increase in excitatory postsynaptic potential (EPSP) amplitude in the postsynaptic cell (Dudek & Bear, 1992). This is due to N-methyl-D-aspartate (NMDA) receptor activation increasing calcium ion levels in the postsynaptic neuron, which with high frequency stimulation results in the addition of more α-amino-3-hydroxy-5-methyl-4-isoxazolepropionic acid receptor (AMPA) receptors to the synapse as well as synapse strengthening genetic changes, and with low frequency stimulation results in removal of AMPA receptors and genetic synapse weakening. For LTP, this change makes it more likely that the postsynaptic neuron fires shortly after the presynaptic neuron, causing an increase in correlation. Paired pulse facilitation refers to the way in which if a neuron is stimulated twice in a row, leftover calcium ions in the synapse from the first action potential increases probability of neurotransmitter release from the second stimulation, resulting in temporary facilitation (Santschi & Stanton, 2003). Unlike LTP, this effect is short term, as after a longer period of time the ions would be cleared, while LTP’s effects can last from thirty minutes to several hours and even for months in some cases (Kumar, 2011). Connectivity changes such as this, especially when long term, can alter the way that the entire local network interacts, and may be responsible for the success of many neuromodulatory therapies, making the ability to induce this type of change with tFUS highly desirable.

tFUS may in fact have these capabilities while remaining entirely non-invasive, unlike electrical stimulation. It has been shown to be capable of targeting neurons on a circuit level safely, as well as affecting large scale connectivity between brain regions in fMRI studies, causing the targeted brain region to either become much more or much less correlated with other brain regions for several hours past the stimulation time (Dallapiazza et al., 2018; Sanguinetti et al., 2020; Verhagen et al., 2019). Previous studies from our group have shown that tFUS is capable of inducing LTD through stimulation at 50 Hz in the hippocampus targeting the medial perforant path (Yu et al., 2019). However, this same frequency in electrical stimulation induces LTP instead, demonstrating that there is a difference in mechanism between tFUS and electrical stimulation that directly affects plasticity changes. This indicates the need for further research into the relationship between ultrasound stimulation frequency and plasticity changes, as it is not parallel to the paradigms studied for electrical stimulation. There is also a need to explore how tFUS affects connectivity within the small circuits in a targeted region, as past studies have also focused more on large scale connectivity between brain regions or directly neuron to neuron. Understanding this level of connectivity modulation as well as the effect that induced changes have on the functionality of the targeted region across different stimulation parameters is critical to understanding the ability of ultrasound as a therapeutic neuromodulator, as many phenomena arise out of such circuits. To analyze connectivity, one helpful tool is measuring the correlation between neurons.

Correlation measures whether one neuron firing is associated with a greater chance of the other one firing within a short time as well (Cohen & Kohn, 2011; Cutts & Eglen, 2014; Tchumatchenko et al., 2011). LTP, by strengthening the synapse between neurons and making it more likely for one to fire after the other, would increase correlation between directly connected neurons. Plasticity changes of the type we hope to demonstrate possible from tFUS would thus be visible through measurement of correlation between targeted neurons. However, there is not only correlation between directly pre- and post-synaptic neurons. A neuron being directly pre-synaptic to the other will result in correlation, but two neurons having the same pre-synaptic neuron will also result in correlation (Tchumatchenko et al., 2011). Pairwise correlation analysis thus helps show how neurons broadly are firing with relation to each other even if not directly connected and shows whether plasticity has occurred somewhere, although it may be hard to completely pick apart the local circuit structure, making it possible for us to determine what changes tFUS may be able to induce.

Here we present an investigation of tFUS’s neuromodulatory effect on neuronal firing and correlation in the S1HL region of the rat brain in response to peripheral electrical stimulation at a unilateral limb. We present evidence that the delivery frequency of pulsed sonication, i.e., sonication repetition frequency (SRF), is important to the induced correlation changes, and that similar to electrical stimulation, low SRFs cause an overall decrease in correlation while high SRFs cause an increase in correlation, but at a shifted set of frequencies compared to electrical stimulation. We show that ultrasound at all of these frequencies causes an increase in firing in response to stimulation, but that the amount of change is related to the correlation changes induced. Finally, we study the different correlation effects found across and within layers of the cortex recorded from, and present evidence that correlation changes induced may vary based on cell morphology variation across layers. Overall, we found that ultrasound is capable of causing overall local network decorrelation or correlation at different frequencies as well as excitation on an individual neuronal level.

## Methods

### Setup and Experimental Design

Thirty-two adult male Wistar rats were studied according to a protocol approved by the Institutional Animal Care and Use Committee of Carnegie Mellon University. Subjects were adults with weight range 400-600 g. In each animal, craniotomy was performed to create a 2 mm burr hole allowing the electrode to target the S1HL region of the rat brain (ML: −3 mm, AP: −0.84 mm, depth 1 mm) and neural recordings were taken with a 32-channel electrode inserted at a 30° degree incidence angle (Figure 1A,1B). The electrode used was a 10 mm single shank electrode with three columns of ten channels on each side column and twelve in the middle column, with electrodes arranged to be 50 microns away from each other vertically and horizontally (A1×32-Poly3-10mm-50-177, Neuronexus, Ann Arbor, MI, USA). A TENS unit electrical stimulation device was attached to the leg with conductive gel placed at the dual electrode contact points for peripheral stimulation (MLADDF30, ADInstruments Inc., Colorado Springs CO, USA). Needle EMG electrodes were inserted in the hind limb to co-register peripheral stimulation reference to the brain recordings. The ultrasound was also delivered at a 30° degree incidence angle from the opposite angle to the recording electrode so that the recording was done from the area being targeted by the ultrasound with peak intensity, with profile shown in Figure 1C. Peripheral electrical stimulation was delivered with a duration of 65 μs in pulses at 0.2 Hz (every 5 seconds) with intensity ranging from 5-11 mA, decided based upon the minimum amount that elicited a visible leg twitch. For surgery, each animal was anesthetized with isoflurane delivered at 3% with a flow rate of 0.4 L/m through a nose cone. After the surgery, isoflurane was reduced to 2%, and a minimum of 30 minutes was given before recordings for anesthesia levels to induce a constant depth of sedation. In each session, the pulses of peripheral electrical stimulation were first delivered to the right hind limb, with stimulation once every five seconds for thirty minutes. Following this, ultrasound stimulation was delivered for five minutes. After ultrasound stimulation, pulses of peripheral electrical stimulation were delivered for sixty more minutes. This experimental paradigm is illustrated in Figure 2A and 2B. Ultrasound stimulation was delivered at 5 different SRFs, i.e., 10 Hz, 50 Hz, 75 Hz, 100 Hz, and 125 Hz. 50 Hz stimulation was observed by our group to induce LTD in the rat hippocampus (Niu et al., 2022a), so this range of frequencies was chosen to further explore the parameter space and discover whether inducing LTP as well may be possible. In each animal, two ultrasound sessions and a sham session were performed in a random order before euthanasia. Fifteen sessions of 50 Hz and 100 Hz were performed, and ten sessions of 10 Hz, 75 Hz, and 125 Hz were performed, with thirty-two animals used total. A sham condition with the ultrasound directed at a different region of the brain at the anterior of the skull was used.

**Figure 1:**
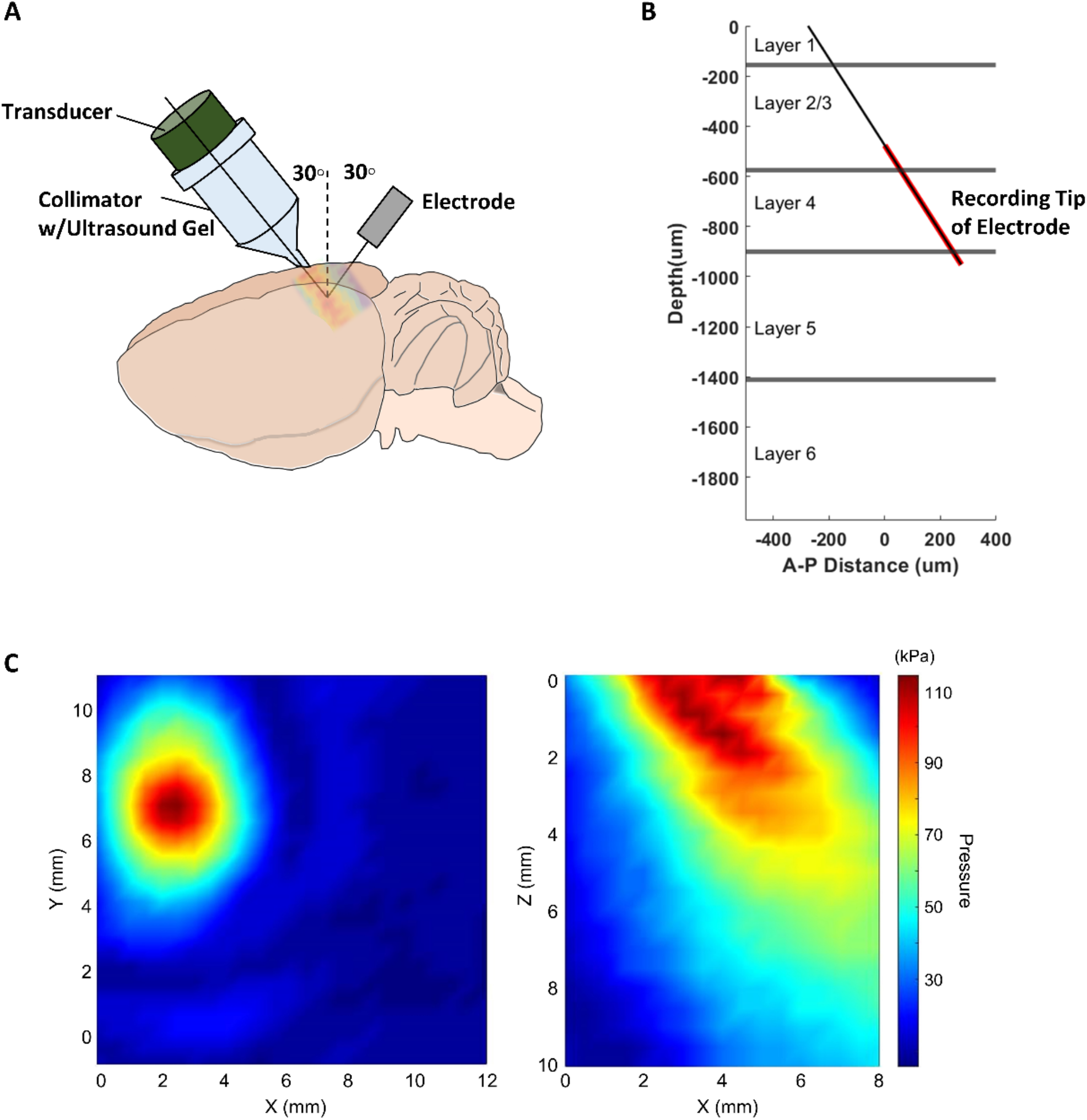
Illustration of ultrasound delivery and electrode insertion. **(A)** Illustration of the positioning of the ultrasound collimator relative to the rat and the electrode, shown in grey. **(B)** Diagram of the electrode insertion depth relative to the estimated depth of each layer of the rat somatosensory cortex. **(C)** Hydrophone pressure scan resulting from the ultrasound parameters used and angled incidence.

**Figure 2.**
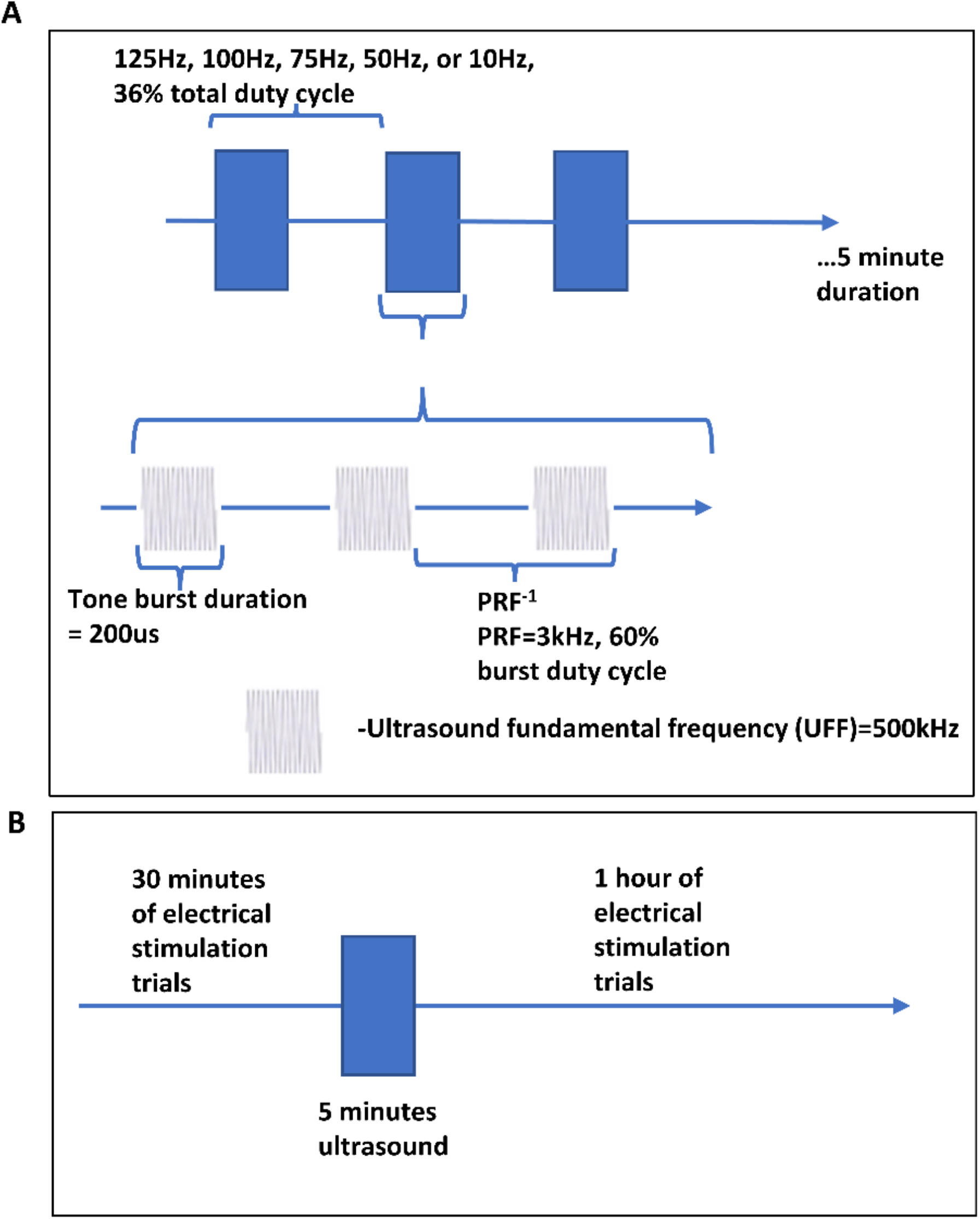
Diagram of Experimental Protocol. **(A)** Diagram of ultrasound waveform and pulsing parameters used during the 5-minute stimulation duration. The sinusoidal blocks represent the fundamental frequency of ultrasound, which is delivered in pulses shown on the middle level, and the blue blocks represent a series of those pulses. These blocks are also delivered in a pulsed mode at a frequency titled the ‘Sonication Repetition Frequency’. SRF was the parameter focused on in this study. **(B)** Diagram of the overall experimental timeline, with the line representing recording time with electrical stimulation being delivered, and the blue block representing the delivery of ultrasound stimulation.

### Ultrasound Setup

Ultrasound was delivered in pulses using a single element focused transducer with a concave surface (AT31529, Blatek Industries, Inc., USA), with an outer diameter of 25.4 mm, fundamental frequency f_0_ of 0.5 MHz, and a nominal focal distance of 38 mm. The transducer was attached to a collimator filled with ultrasound gel which was pointed towards the targeted brain region (Figure 1A). The collimator used was 3D printed with VeroClear material to be a size matching the focal length of the transducer and with an outlet greater than or equal to the ultrasound wavelength (~3 mm in soft tissue), specifically with elliptical area 25.6 mm^2^ (major axis 6.8 mm, minor axis 5 mm). Three function generators were used, one double-channel waveform generator (33612A, Keysight Technologies, Inc., USA) and two single-channel waveform generators (33220A, Keysight Technologies, Inc., USA). Together these generated the fundamental frequency f_0_ of 500 kHz, the PRF of 3 kHz, and the SRF of 10-125 Hz as portrayed in Figure 2A. A power amplifier was used on the low voltage ultrasound waveform before delivery (BBS0D3FHM, Empower RF Systems, Inc., USA). The tone burst duration was 200 μs. Ultrasound is delivered with a burst duty cycle of 60%. Stimulation is also given at 60% duty cycle, resulting in a total duty cycle of 36%.

### Electrophysiological Preprocessing and Analysis

For spike analysis, recordings were bandpass filtered between 300 Hz and 6 kHz. Spike sorting was performed using Kilosort 2.0 (Pachitariu et al., 2016). Phy2, a manual spike sorting GUI was then used to check the output of Kilosort for errors, and merge or split any discovered clusters that were sorted incorrectly based off of inter-spike-interval histograms as well as clustering of PCA derived waveform properties. Recordings were bandpass filtered from 1-300 Hz for LFP continuous data and AC line noise at 60 Hz was filtered as well. Further analysis was completed using custom code in MATLAB (R2019a, MathWorks, USA). Peak detection was used on the EMG recordings using a needle electrode recording from the limb to find the stimulus onset times. Spike times for recorded neurons were then sorted into their peripheral electrical stimulation trials, with approximately 360 trials before ultrasound and 720 trials afterwards. Peri-stimulus time histograms (PSTH) were created for each neuron before and after ultrasound. The difference between average number of spikes elicited by electrical stimulation in the 0.5 s following stimulation and the average number of spikes at baseline was recorded for each neuron for 120 trials (ten minutes) before and after ultrasound stimulation. The average value for every ten minutes of recording was also plotted over the entire recording time to ensure stability.

To analyze correlation, the Spike Time Tiling Coefficient (STTC) was used (Cutts & Eglen, 2014). This metric is a way of measuring whether one neuron spiking increases the probability that the other neuron will also spike within a short time frame, while compensating for differences in firing rates between neurons which can otherwise bias correlation metrics. The STTC is calculated through equation (1).

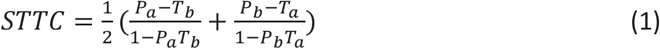

Where T_a_ and T_b_ represent the proportion of recording time which is within some set Δt from a spike of neuron A and B, respectively. P_a_ represents the proportion of spikes from neuron A which are within Δt from any spike of neuron B, and P_b_ represents the proportion of spikes from neuron B which are within Δt from any spike from neuron A. This formula is based on the assumption that the proportion of spikes from neuron A within Δt of a spike of neuron B by chance is equal to the proportion of time in the recording within Δt of a spike from B to the total time. Following this assumption, P_a_ = T_b_ if there are only the expected number of spikes within Δt given the firing rate, and P_a_ - T_b_ is positive if there is a higher correlation. The other terms in the equation are used to give the coefficient the property of symmetry and normalization such that the coefficient ranges from −1 to 1. Δt was chosen to be 50 ms, a value within the range commonly used in literature (Bair et al., 2001; Cohen & Kohn, 2011; Cutts & Eglen, 2014). Like in the previous analyses, correlation was measured in the 120 trials (ten minutes) before and after ultrasound stimulation. Although some measured correlation will be due to the electrical stimulus, it will be common before and after ultrasound stimulation and thus negated. The value was also plotted for every ten-minute time bin across the recordings to ensure stability.

The depth along the recording probe of each neuron was used to sort the neurons into their estimated cortical layer. Instantaneous firing rate was measured to check whether highest firing rates were found in L5 as expected from literature(Narayanan et al., 2017a; Ryu et al., 2019). Correlation in each layer were then compared using the same process and metric as stated previously.

### Statistical Analysis

Data from neuronal spiking and correlation were tested for normality using the Shapiro-Wilk test, and the distributions were found to be not normal. Following this, group analysis of significance was performed using the Kruskal-Wallis H test and significance characterized by *post hoc* Wilcoxon tests with the Bonferroni correction for multiple comparisons (\**p* < 0.003, ** *p* < 0.00139 for layer correlation analysis). Some sessions were excluded from analysis due to a reduction in quality mid-session, likely due to the local surgical and electrode insertion disruption resulting in the death of neurons, which made spike sorting for parts of the session impossible.

## Results

### Spiking Response

Electrical stimulation to the hind limb prompting a neural response in the somatosensory cortex was used to characterize functionality of the somatosensory cortex. After electrical stimulation, there is a burst of activity in the neurons followed by a quiet period and then a return to normal levels (Figure 3A). Following ultrasound stimulation, all frequencies on average resulted in an increase in firing in response to electrical stimulation (Figure 3B-D) that was statistically significantly different compared to the sham condition (*p* < 1e-13). In sham sessions, either no change or a decrease in response was observed. Increased firing rates generally lasted for 10-20 minutes, although occasionally it lasted for the remainder of the session (Figure 3C). The 10 Hz condition showed the highest increase, significantly (*p* < 0.002) more than those in 125 Hz and 75 Hz conditions. 75 Hz showed the least change, although only significantly distinct from sham and 10 Hz. This appears to indicate that our stimulation paradigm was overall excitatory in nature due to the use of PRF at 3,000 Hz.

**Figure 3:**
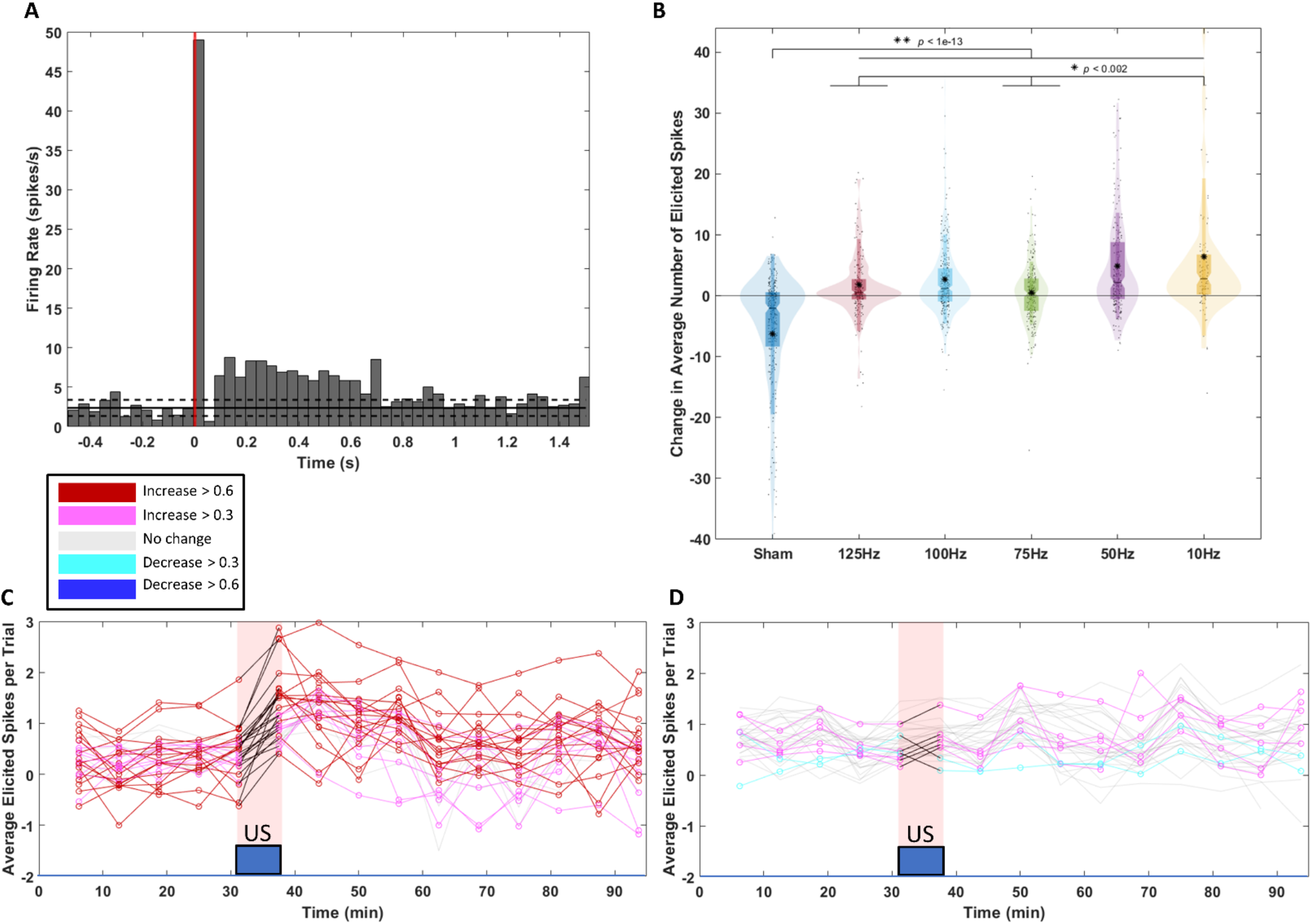
Spiking Response Change following Ultrasound Stimulation. **(A)** Characteristic peri-stimulus time histogram of the response to hind limb electrical stimulation of a neuron’s firing over trials. **(B)** Violin plot of the changes in spiking response to hind limb electrical stimulation following tFUS stimulation at S1 in each experimental condition. Each black dot represents a neuron from a recording under the labeled SRF. The black asterisk marks show the mean of each group. The darker color is a traditional violin plot, with the ends of the boxes marking the 1^st^ and 3^rd^ quartile, and the middle black line showing the median. The notches show 95% confidence intervals on the median. The ends of the narrow rectangle show the mean plus and minus one standard deviation. The lighter color shadow over the data shows the probability distribution of the data collected. The ends of that distribution mark the 1^st^ and 99^th^ percentile of the data. **(C-D)** Characteristic plots of the average spiking response over time during an experiment for a true ultrasound condition in (C), and in a sham condition in (D). Each line represents a neuron in the recording, with the color of the line representing the degree of change over ultrasound stimulation. The duration of ultrasound stimulation is shown by the blue box. Each point in the line is the average over five minutes of the number of spikes different from baseline in the half second timebin following hind limb electrical stimulation for a neuron.

### Correlation Change

Following our group’s previous work in the hippocampus examining the ability of ultrasound to induce plasticity changes(Niu et al., 2022b), we next examined whether ultrasound was capable of causing correlation changes in S1. Examining the changes in correlation induced between every pair of neurons in a session, it was observed that every frequency resulted in a significant average correlation change besides 10 Hz (*p* < 1e-4) (Figure 4G). The 100 Hz condition was significantly higher than all other conditions (*p* < 1e-7), while the 50 Hz condition was significantly lower than all except the 10 Hz condition (*p* < 1e-5). This interestingly appears to align with the previous hippocampus result of 50Hz causing LTD (Niu et al., 2022b)Here, we observe that in response to 50 Hz ultrasound stimulation, on average, the network undergoes a decrease in correlation. In about half of the cases these correlation changes appear to last for the remainder of the session time, although in others the change only lasts for 10-20 minutes much like the firing rate changes (Figure 4A-C). Interestingly, in most sessions at all conditions there are a mix of positive and negative correlation changes beyond. To look further into this, we observed which correlation changes appeared to be significant within each recording. Significant correlation changes were defined as those with a change greater than 0.1, which was the average standard deviation of a baseline recording’s correlation over time, and which were also greater than the standard deviation of the baseline correlation for the specific recording. When plotting these correlation changes both over time and in a representation of the neurons’ approximate location relative to the recording probe, the difference in effect between 100 Hz and 50 Hz conditions is clearer. Although large changes in correlation may be induced in both directions between individual pairs, the overall effect is an increase and decrease in correlation for 100 Hz and 50 Hz respectively (Figure 4A-F). 125 Hz and 75 Hz appear to have effects similar to the 100 Hz, largely resulting in a network-wide increase in correlation, while 10 Hz is more similar to the 50 Hz condition, resulting in an overall decrease in correlation (Figure 4G). When examining the percent of significant correlation changes the frequencies all appear to have similar effect sizes. However, breaking them down into positive and negative changes, 125 Hz, 100 Hz, and 75 Hz all show more positive changes than negative, while 50 Hz and 10 Hz show more negative changes than positive (Figure 4H). This indicates that there may be a specific threshold between 50 Hz and 75 Hz where the overall effect changes from positive to negative. The degree of correlation change even within a frequency however appears to vary significantly. Examining the correlation changes in spatial form, it was also observed that there appear to be a few neurons which are responsible for the majority of the significant correlation changes (Figure 4D-F). This may indicate a particular effect on those neurons, among those recorded.

**Figure 4:**
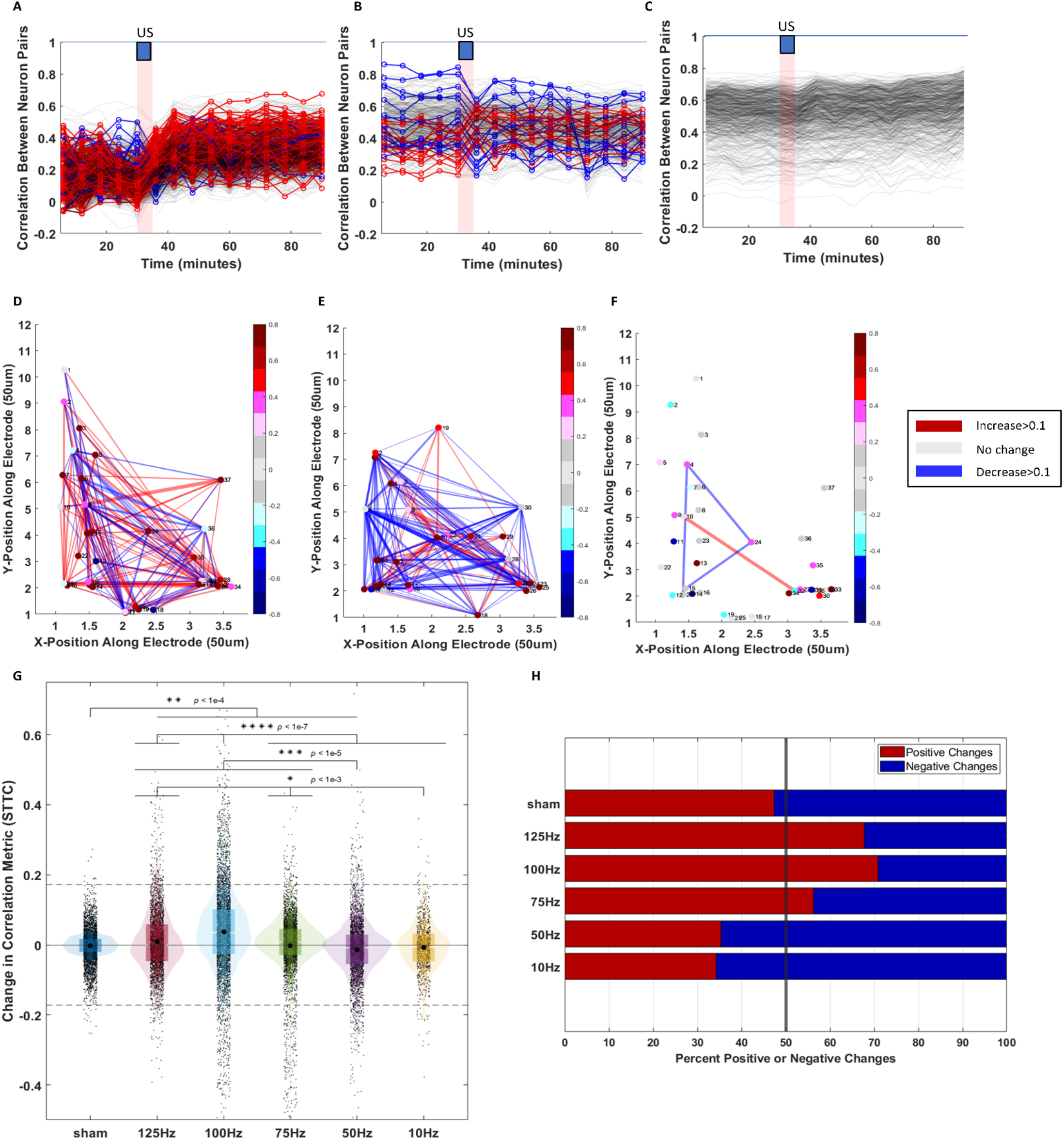
Correlation Change following Ultrasound Stimulation. **(A-C)** Characteristic plots of correlation over the time of an experiment at 100 Hz SRF (A), 50 Hz SRF (B), and in the sham condition (C). The STTC was measured over five-minute time intervals and each line represents a pair of neurons in the recording, with the color of the line representing the direction and degree of change after ultrasound stimulation. **(D-F)** Graphical plots showing the change in correlation between neurons with each point representing a neuron in the recording, and lines between neurons representing significant correlation changes following ultrasound. 100 Hz SRF is shown in (D), 50 Hz SRF in (E), and a sham condition in (F). The color of the line represents the direction of correlation change, and the thickness of the line represents the scaled degree of change, with a thicker line showing more change in the relevant direction. The location of each neuron is approximated by the channel of the recording electrode with the highest amplitude. The color of the node for each neuron represents the change in firing response to hind limb electrical stimulation in that neuron. **(G)** A violin plot plotting correlation change between pairs of neurons in each experimental condition. The dashed line marks three standard deviations outside of the mean of the sham group. The sham condition shows minimal correlation changes, with the group tightly clustered around 0. By contrast, all other groups except 10 Hz show changes in correlation between many neurons. 75 Hz and above show generally positive correlation changes, while 50Hz and below show generally negative correlation changes. **(H)** A bar plot showing the ratio of positive and negative correlation changes at each SRF, with a positive change showed in red and a negative one in blue.

Neurons were then sorted into their putative layer position, and correlation within the layers at 100 Hz and 50 Hz was analyzed. Overall, among 100 Hz, 50 Hz, and sham, the level differences from the previous correlation analysis remained (Figure 5B-D). Within a condition, interesting differences among layers exist. At 100 Hz, L3 stands out as having more increases in correlation than L4 or L5, although it is only significantly higher than L5 with *p* < 1e-7. L4 has a higher variance than the other layers, although this may be due to it also having a higher number of neurons analyzed. At 50 Hz there is no significant difference between the layers, although once again L4 shows the highest variance in response. The sham condition interestingly shows a decrease in level correlation in L5, however it is not a significant difference. Overall, these different distributions of response may indicate that morphology differences between neurons in the different layers of cortex may have intrinsically different responses to ultrasound stimulation.

**Figure 5:**
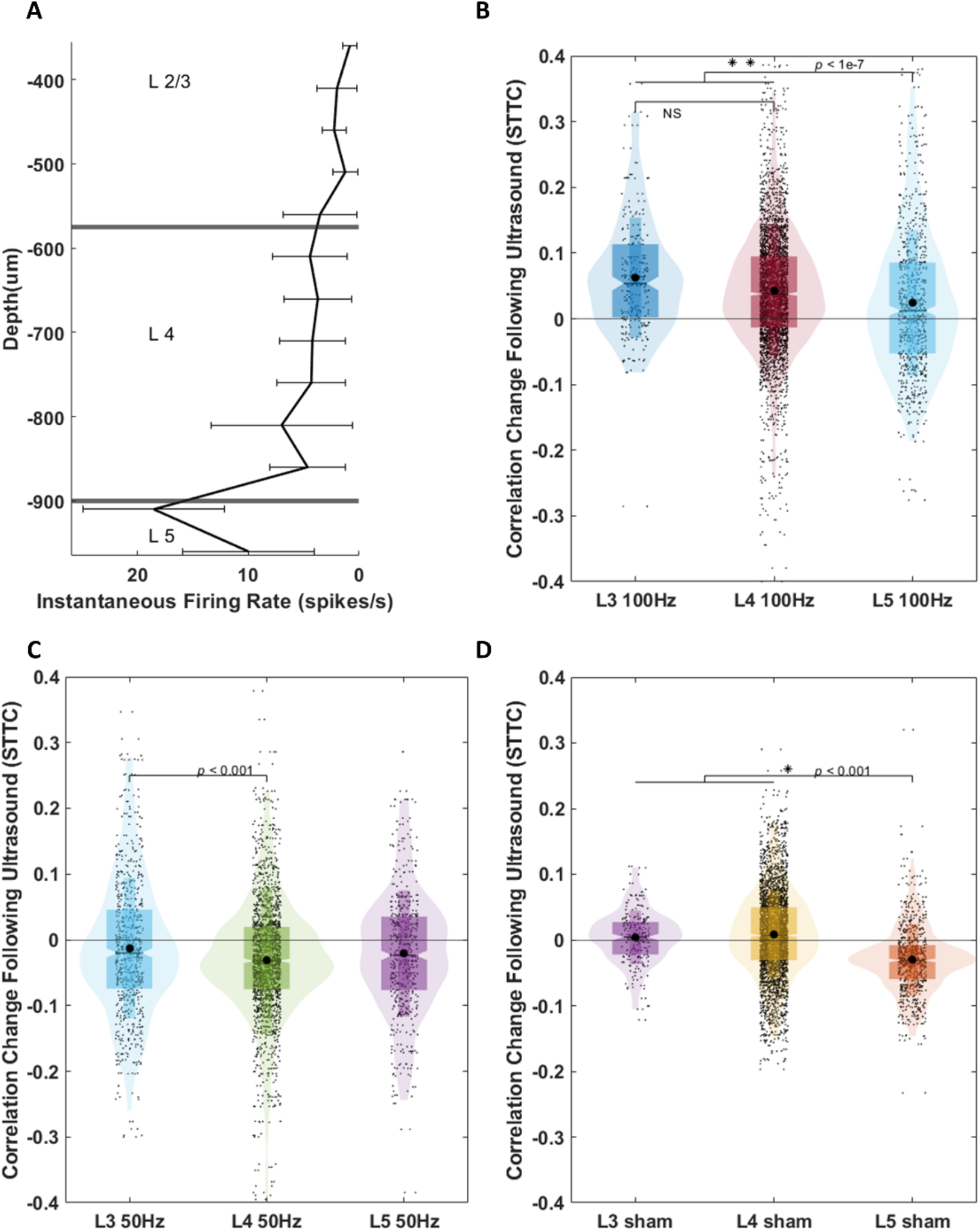
Layer Analysis of Correlation Change. **(A)** A plot showing the instantaneous firing rate at each depth in the cortex, assisting in demarcating the boundaries between layers. **(B-D)** Violin plots with each point plotting correlation change between neuron pairs following ultrasound. In (B), the neuron pairs within Layer 2/3, Layer 4, and Layer 5 are shown for the 100Hz condition. (C) the 50Hz condition, and (D) the sham condition.

## Discussion

### Excitatory effects in response to electrical stimulation following ultrasound

The neuronal firing response in S1 to peripheral electrical hind-limb stimulation was measured before and after ultrasound at five different SRFs and one sham condition. In all of the ultrasound conditions, neurons were broadly found to increase their spiking activities when they responded to electrical stimulation relative to baseline after ultrasound stimulation. This appears to indicate an overall excitatory effect of ultrasound stimulation. Ultrasound has been shown to induce lasting excitation *in vitro* as well as in EEG (Clennell et al., 2021; H. C. Kim et al., 2021), but to our knowledge this is the first time this effect has been shown in intracortical single unit recordings. Within typical stimulation paradigms, it has been generally found that a shorter sonication duration combined with a high duty cycle (≤500ms, ≥30%) results in excitation while a longer duration and lower duty cycle (≥1minute, ≤10%) results in suppression (H. C. Kim et al., 2021; Yoon et al., 2019). In our paradigm, we deliver typical ultrasound stimulation in pulses to mimic the delivery of electrical stimulation in pulses that has been used to induce LTP and LTD. Our total duty cycle (36%) and tone burst duration of 200 μs line up with the excitatory range previously described. We find here that delivering this stimulation in pulses does in fact create an excitatory effect, even with the five-minute overall duration. We also find that the duration of this effect lasts beyond the stimulus, enduring for at least 10-20 minutes and sometimes past an hour after stimulus (Figure 3C). In the sham condition, although the majority of neurons showed no changes, some showed a decrease after ultrasound. Decreases are also seen in some true ultrasound sessions, which may demonstrate a baseline random variation in response, but there are clearly more large decreases in the sham condition. This may be due to acclimation to the stimulus, although we cannot rule out it being the effect of the sham ultrasound (off target) from a distance. Although all true ultrasound conditions show an increase, there are some significant level variations among them. 10 Hz notably shows an increase over 125 Hz and 75 Hz. Overall, 75 Hz shows the least change despite still being significantly different from sham. The variation may show that the excitation is frequency dependent in a way related to SRF, and potentially the effects on connectivity observed. It is notable that despite the overall excitatory effect on average, not all neurons respond the same way. This may represent different ultrasound effects on subtypes of neurons, or possibly a variation in ultrasound effect across space due to slight variations in physical structure. Although a clear trend is not obvious, it appears that this excitation does vary with SRF.

### Correlation changes following ultrasound stimulation

Correlation between neuron pairs were measured in ten-minute time-bins over the experiment, before and after ultrasound in all conditions. Ten minutes was chosen since it was the minimum time that changes appeared to exist over, without gradual decrease minimizing the visibility of the initial effect in some sessions. Four of the ultrasound conditions showed significant average correlation changes compared to sham. Looking at individual sessions (Figure 4 A-C), a clearly visible change in the correlation between neurons from relatively stable baseline states is present in true ultrasound conditions compared to sham. When looking at the effect on individual neurons (Figure 4 D-F) it appears that there are specific neurons which have strong correlation changes with a large number of neurons, much more than others. We hypothesize that these represent the neurons which ultrasound stimulation succeeded in stimulating in a manner causing synaptic plasticity changes, and that more minor correlation changes in other neurons may be resulting from downstream impacts of these bigger correlation changes. Like in the firing response case, this may be due to either neuronal sub-type variation, or other more general structural variation local to the neuron such as astrocytic structures (Oh et al., 2019). Changes are notably bimodal in nature at all SRFs, but when looking at the ratio between positive and negative changes at different SRFs, we observed that 75 Hz, 100 Hz, and 125 Hz appear to broadly cause increases in correlation on average, while 50 Hz and 10 Hz cause decreases on average. This is significant because low frequencies causing decreases in correlation and high frequencies causing increases is a pattern observed in the electrical induction of synaptic changes through LTP and LTD. However, it is unclear if that is the (sole) mechanism at work. Correlation changes sometimes lasted for up to an hour after ultrasound stimulation, which may suggest LTP or LTD, but other times the correlation changes only persisted for 10-20 minutes following ultrasound. Closer examination and large numbers of animal sessions may be required to fully parse out the mechanism involved here. The bimodal nature also requires further investigation. If LTP and LTD are at play, it may be that the imprecise nature of ultrasound’s stimulation of neuronal firing is the cause of there being both increases and decreases. With electrical stimulation, each stimulus causes the local neurons to fire by directly affecting voltage and causing an action potential. There are many theorized mechanisms for ultrasound’s ability to cause neuronal firing including intramembrane cavitation, radiation pressure, thermal energy, and mechanosensitive ion channels including in astrocytes, but in all of them the timescale of effect is much slower then electrical stimulation (H. Kim et al., 2014; Naor et al., 2016). This means that changes in SRF may only imprecisely increase the rate at which a neuron is induced to fire, while with electrical stimulation, firing rate is directly created by the stimulation frequency. At 100 Hz, ultrasound might consistently cause the neurons to fire at a frequency allowing synaptic strength increases, while at lower frequencies the effect may be more mixed. To examine this hypothesis, in depth study of the firing rate induced by different SRFs would be helpful. It is also possible that there are varying responses due to the timing of sonication relative to the baseline phase of activity the neuron is in. Regardless, we have shown that ultrasound is capable of having significant effects on the correlation between neurons network wide. Correlation change does not appear directly tied to firing response changes, as neuron with both correlation increases and decreases showed excitation following tFUS. However, it was observed that the 75 Hz condition resulted in both the least average change in firing response and correlation change. Neurons with large correlation changes generally also had significant firing response changes. This appears to show that the ultrasound effect causing both of these changes is somewhat linked.

When the neurons were separated into their putative different layers of cortex, variations in the correlation change effect were also found. It is known that different subtypes of neurons exist within the different layers of cortex. For example, L3 mostly contains regular pyramidal cells, while L4 contains pyramidal cells, spiny stellates, and star pyramids and L5 contains slender and thick tufted pyramids (Narayanan et al., 2017b). All of these have different shapes and sizes and may be interacted with a mechanical wave differently, resulting in the slightly different correlation changes observed.

### Future Directions and Potential Confounds

This paradigm of ultrasound stimulation delivered in pulses of pulses of the fundamental frequency as shown in Figure 2 is not well explored with regards to excitatory or inhibitory effect, as opposed to merely a single layer of pulses. Usually, ultrasound is either delivered continuously or just in single level pulses of the fundamental frequency. Since pulses of ultrasound are what have generally been studied to have modulatory effects, we used pulses of this stimulation to mimic pulses of electrical stimulation. If these pulses could successfully stimulate neurons to fire at the frequencies necessary to induce plasticity effects such as LTP or LTD, we hypothesized that we would see correlation changes as a result. Our findings confirm these changes, supporting our hypothesis, but whether this is the only mechanism at work remains unclear. We also hypothesized that the parameters used in pulses would maintain their previously seen success in neurostimulation. Since excitation was found to occur, this hypothesis was generally supported, but it was not shown in this study whether the parameters generally have exactly the same effects within this stimulation paradigm as in the more common one. This study focused on SRF as a parameter, but there remain many questions as to how other parameters such as PRF, UFF, and both burst and total duty cycle affect the results observed in this study. Previous studies have shown that parameters such as PRF can induce cell-type specific effects, making it likely that varying PRF could result in significantly different network effects than were observed here (Yu, Niu, et al., 2021). Future studies must examine the large parameters space that ultrasound stimulation has in order to determine the variety of effects possible.

Ultrasound neuromodulation also comes with certain potential confounds due to its nature as vibration. In particular, it has been noted that it is possible for ultrasound to have effects through an indirect auditory activation pathway (Kamimura et al., 2020; Sato et al., 2018). To account for this possibility, we use a sham condition in which ultrasound is also delivered, but to a different region of the brain the same distance from the auditory cortex. Through this control, any indirect auditory effects should also exist in the sham condition, allowing us to separate more direct tFUS effects from the auditory mechanism. Another concern raised in ultrasound stimulation specifically when recording from an electrode is whether the electrode tip itself is vibrating and causing local effects through its movement. Previous studies have found that electrode displacement becomes significant past the threshold of a pressure of 131kPA, which our stimulation paradigm does not reach (M. G. Kim et al., 2021). This suggests that electrode vibration is not a concern in this research.

Besides ultrasound effects, the effects of anesthesia are another potential confound to the results found here. In this study, isoflurane through inhalation is used to keep the animal subjects anesthetized. Anesthesia is known to be a major confound in neuromodulation studies, given that it is known to affect spontaneous firing rate, spiking patterns, and result in overall depression of neural activity. Some theories of how anesthetics work involve mechanisms that interact with theorized mechanisms of tFUS, specifically in interaction with membrane excitability (Kamimura et al., 2020). These confounds could potentially affect several parts of this study, as anesthesia level directly modulates each of the metrics examined here. To reduce these confounds, anesthesia is kept at as low as possible a level while keeping the animal sedated enough to prevent animal movements which would break the recording electrode. Additionally, the level is kept stable throughout while monitoring animal vitals, and an acclimation period to the low anesthesia level is given before recordings are taken, to allow any effects to become stable. Despite these efforts, awake studies may be required to fully confirm whether the same level of effect is seen in awake subjects, as would be important if tFUS were to be used as a therapy for neurological disorders.

The results in this study further confirm the ability of ultrasound to affect neuronal correlations in a parameter dependent manner. Certain ultrasound SRFs resulted in overall correlation increase, while others resulted in overall correlation decrease. The time scale over which these changes last varied in experiments, but consistently persisted beyond the stimulus duration, further confirming the ability of ultrasound to have a lasting effect beyond the duration of stimulation. We also observed the ability of ultrasound to modulate neuronal firing through intracranial recordings, with some dependences on the SRF. These demonstrations of the wide capabilities of tFUS and its flexibility in effect depending on parameters used are promising for its future as a neuromodulation therapy.

### Conclusions

We observed that ultrasound can induce excitation in individual neurons responding to peripheral stimulation, and that the level of excitation is frequency dependent. Extending previous results, we found that ultrasound induces pairwise correlation changes between targeted neurons. Sustained decorrelation was found to be generally induced at low frequencies (< 75 Hz) including the frequency at which synaptic depression was found previously, 50 Hz, and sustained potentiation was found at higher frequencies, in particular 100 Hz. This demonstrates that ultrasound has the capability to cause plasticity changes in both directions by adjusting the parameters used.

## Acknowledgment

This work was supported in part by NIH grants MH114233, EB029354, NS124564, AT009263, EB021027,and NS096761.

## Data availability

All data that support the findings of this study are included within the article.

